# Two-State Effective Connectivity Predicts the Effect of Anterior Cingulate Cortex Excitatory/Inhibitory Balance on Posterior Cingulate Cortex

**DOI:** 10.64898/2026.09.01.748001

**Authors:** Abdoreza Asadpour, Aanya Malaviya, James Cook, Alan George, Piyali Bhattacharya, Ifigeneia Manitsa, Martin P Wilson, Maria R Dauvermann

**Affiliations:** Sussex Neuroscience, School of Life Sciences, University of Sussex, Brighton, UK; Institute for Mental Health, School of Psychology, University of Birmingham, Birmingham, UK; Centre for Human Brain Health, School of Psychology, University of Birmingham, Birmingham, UK

**Keywords:** Excitatory/inhibitory balance, Glutamate+Glutamine/GABA ratio, posterior cingulate cortex, Dynamic Causal Modelling, resting-state fMRI, default-mode network, psychotic experiences, Magnetic Resonance Spectroscopy

## Abstract

**Aims:** Excitatory/inhibitory imbalance - the equilibrium between glutamatergic excitation and γ-Aminobutyric acid (GABA)ergic inhibition - is implicated in psychosis and may precede psychosis onset.

**Methods:** To investigate whether excitatory/inhibitory balance relates to subclinical psychotic experiences, anxiety and depression, we developed a network-level measure integrating Magnetic Resonance Spectroscopy Glutamate/GABA ratios with effective connectivity during resting-state fMRI. In 20 healthy young adults (aged between 18-32; 50% female), we first measured excitatory/inhibitory balance, operationalised as Glutamate/GABA and Glutamate+Glutamine(Glx)/GABA ratio, in the dorsal anterior cingulate cortex. Second, we applied Two-state Dynamic Causal Modelling to assess effective connectivity of the default-mode network, using these ratios as empirical priors to determine how they translate into network connectivity. Third, we ran Bayesian Model Selection at the model and family levels to identify the optimal models.

**Results:** The two-state model with Glx/GABA ratio as direct input into the posterior cingulate cortex best explained the data. This best-fitting model incorporated the neurometabolite ratio with Glutamate and GABA and used two-state connectivity.

**Conclusions:** Excitatory/inhibitory balance underlies resting-state effective connectivity, with the Glx/GABA ratio shaping connectivity networks. This pathway highlights excitatory/inhibitory balance as a predictor of two-state brain networks, potentially aiding identification of high-risk individuals and informing new interventions.

## Introduction

Psychotic disorders, such as schizophrenia, affect approximately less than 1% of adults in England^1^ with an estimated total monetary cost of £11.8 billion for schizophrenia and psychoses in England^2^. Psychotic disorders can have devastating effects for individuals and their families. Early treatment or preventative treatments could help improve outcomes and reduce societal and personal costs. Some individuals are at increased risk of transitioning to a psychotic disorder (e.g., clinical high risk), however identifying these people is difficult because current diagnostic methods are hampered by the absence of accurate objective biomarkers, leading to delays and inaccuracies in diagnosis. Notably, there is some evidence that individuals who are at high risk of developing psychosis show alterations in brain function and networks, and this may predict later transition to a full psychotic disorder^3,4^.

A growing body of evidence shows that the balance between excitatory (mainly, Glutamate) and inhibitory (mainly γ-aminobutyric acid (GABA)) regulation is a core neurobiological feature for plasticity and cognitive functioning^5^ on the brain network level. When this balance is disrupted, both increased and reduced Glutamate and GABA concentrations have been reported across psychotic disorders^6,7^. Post-mortem, pharmacological, and *in vivo* Magnetic Resonance Spectroscopy (MRS) studies^8–10^ have consistently reported alterations in Glutamate or GABA concentrations in the prefrontal cortex across psychosis and schizophrenia^8,10^. Studies find alterations with both increased and reduced levels with a trend towards increased Glutamate and Glutamate+Glutamine (Glx) levels in the dorsal anterior cingulate cortex (dACC) in early psychosis^11,12^, whereas reduced Glutamate and Glutamate+Glutamine levels in the medial prefrontal cortex (mPFC) in individuals with schizophrenia than controls^10,13^ are reported. Similarly for GABA, findings of both increased and reduced GABA levels in the mPFC at the early stages of psychosis^14,15^ are observed. Emerging evidence shows a more complex pattern of increased Glutamate and Glx levels with reduced GABA levels in prefrontal regions across different stages of schizophrenia-spectrum disorders^16^, suggesting disruptions of excitatory/inhibitory balance. Despite these advances, a fundamental gap remains. It is not yet understood how neurochemical excitatory/inhibitory imbalance at the regional prefrontal level translates into disrupted large-scale brain network connectivity in individuals at the earlier stages of the psychosis continuum.

It is widely established that resting-state functional and effective connectivity studies reliably have demonstrated altered default-mode network (DMN) connectivity in individuals with psychotic disorders, including schizophrenia^17–20^. Bridging neurometabolic and brain network connectivity, recent evidence suggests that alterations in prefrontal Glutamate and GABA levels are linked to disrupted resting-state connectivity, such as DMN dysconnectivity, in the pathophysiology and prognosis of psychotic disorders, including early psychosis^7,21–25^. Such studies suggest that relationships between Glutamate and GABA in the dACC may be implicated in DMN dysconnectivity^26^. However, these studies are correlational in nature and cannot establish the direction of this relationship between neurochemistry and network connectivity.

Unlike such correlational studies, findings from computational studies modelling resting-state functional connectivity data and excitatory/inhibitory balance in healthy participants suggest that excitatory/inhibitory neurotransmitter homeostasis plays a role in healthy and typical brain network connectivity^27–29^. This hypothesis proposes that excitatory/inhibitory balance between Glutamate and GABA is optimally achieved to ensure typical brain network function in the healthy brain^27^. Yet to date, this theory has not been examined using resting-state connectivity data in individuals with psychotic experiences. Therefore, it remains unknown whether excitatory/inhibitory balance and its associated brain network effects are exacerbated in individuals with psychotic experiences.

To address these gaps, we employ an *in-vivo* and non-invasive multimodal neuroimaging approach combining MRS-derived Glutamate, Glx and GABA metabolites/ratios in the dACC with two-state Dynamic Causal Modelling (DCM) of resting-state fMRI data in individuals with psychotic experiences. In contrast to standard functional connectivity methods, DCM for fMRI enables causal inference by modelling directed influences between brain regions at the neuronal level^30^. In the next step, two-state DCM models were used to measure excitatory and inhibitory contributions to effective connectivity providing a simplified representation of glutamatergic and GABAergic neurotransmission. In the third step, we integrated the Glutamate/GABA and Glx/GABA ratios with the two-state DCM models to examine how Glutamate/GABA ratios in the dACC modulate network properties within the DMN. This approach allows, for the first time, to examine the excitatory/inhibitory neurotransmitter homeostasis in individuals with psychotic experiences. We further evaluate whether these findings are sex specific by comparing females with males. We hypothesised that individuals with greater levels of psychotic experiences and subclinical depressive symptoms will show lower Glutamate/GABA ratios in the dACC when compared to individuals with lower levels of psychotic experiences and subclinical levels of depression. Given the novelty of examining the excitatory/inhibitory neurotransmitter homeostasis theory, we will also explore potential sex differences.

## Methods

### Participants

Thirty-seven healthy young adults (18–35 years; 19 females, 18 males) were recruited via the University of Birmingham, public advertisement, and Call for Participants (callforparticipants.com). None had a clinical diagnosis or treatment history for a health condition, including psychotic disorder, anxiety, or depression (full exclusion criteria in Supplementary Material). All participants provided written informed consent in accordance with local Ethics Committee guidelines.

### Data Collection

#### Demographic data

Standard demographic data and estimated IQ (WASI-II^31^; Vocabulary, Similarities, Block Design, Matrix Reasoning subtests) were collected. Please see the supplementary material for details.

#### Subclinical symptom assessment

Trained, criterion-reliable researchers assessed psychotic experiences (Structured Interview for Psychosis-Risk Syndromes^32^ positive, negative, disorganised, general symptom subscales), subclinical depression (Hamilton Depression Rating Scale^33^), and subclinical anxiety (State-Trait Anxiety Inventory, state and trait subscales) using established interview-based instruments. Please see the supplementary material for details.Trained, criterion-reliable researchers assessed psychotic experiences (Structured Interview for Psychosis-Risk Syndromes^32^ positive, negative, disorganised, general symptom subscales), subclinical depression (Hamilton Depression Rating Scale^33^), and subclinical anxiety (State-Trait Anxiety Inventory^34^, state and trait subscales) using established interview-based instruments. Please see the supplementary material for details.

#### Neuroimaging data acquisition

All participants underwent a single neuroimaging session on a 3T Siemens Prisma system (32-channel head coil) at the Centre for Human Brain Health, University of Birmingham, acquired in the order: T1-weighted, rs-fMRI, single-voxel MRS (semi-LASER and MEGA-PRESS).

### Structural Magnetic Resonance Imaging

A T1-weighted 3D-MPRAGE structural scan was acquired for anatomical reference (1 mm isotropic; full parameters in Supplementary Material).

### Magnetic Resonance Spectroscopy

Single-voxel MRS was acquired in the dACC using semi-LASER³⁵ (Glutamate; 20 x 25 x 20 mm (RL x AP x FH) voxel) and MEGA-PRESS³⁶ (GABA; 30 x 35 x 25 mm (RL x AP x FH) voxel) sequences, with standardised voxel placement in the dACC (Supplemental Figure 1; Supplemental Table 1 on tissue proportion results and full parameters in Supplementary Material).

### Resting-state functional Magnetic Resonance Imaging

Resting-state fMRI was acquired using a multiband GRE-EPI sequence (2.5 mm isotropic; 394 volumes; ∼10 min; full parameters in Supplementary Material).

### Data analysis

#### Neuroimaging data analysis

##### Magnetic Resonance Spectroscopy Data Analysis

Both the semi-LASER and MPRESS MRS data were corrected for single-shot phase and frequency instability using the RATS method^35^ as a preprocessing step. Spectral fitting was performed with ABfit-reg^36^ using a simulated basis-set matched to the MRS acquisition parameters. All MRS processing steps were implemented in the [spant analysis package](https://martin3141.github.io/spant/)^37^ developed for the R programming language.

### Resting-state functional Magnetic Resonance Imaging Data Analysis

#### Pre-processing of rs-fMRI data

Pre-processing and first-level general linear model of the rs-fMRI data was performed using Statistical Parametric Mapping (SPM12; Wellcome Trust Centre for Neuroimaging, London, UK; https://www.fil.ion.ucl.ac.uk/spm) in MATLAB (R2022a; The MathWorks Inc., Natick, MA). Please see the supplementary material for full pipeline details.

### Two-state Dynamic Causal Modelling for fMRI

#### Background to DCM, two-state and stochastic DCM for fMRI

DCM is an established method for assessing inter-regional effective connectivity by modelling experimentally or spontaneously induced changes in neural activity^30^. In contrast to standard functional connectivity methods, which measure correlations between regional time series, DCM enables causal inference by modelling directed influences between brain regions at the neuronal level. It does so by estimating how the rate of change of neural activity in one region influences neural activity in other regions, using coupled differential equations that are linked to predicted BOLD responses via a biophysical haemodynamic forward model^30^.

Unlike the standard (one-state) bilinear DCM, which models each region as a single neuronal population and conflates excitatory and inhibitory contributions, two-state DCM separates each region into excitatory and inhibitory subpopulations^38^, allowing their independent contributions to inter-regional connectivity to be estimated — making it well suited to testing our hypotheses about dACC-DMN excitatory/inhibitory balance. We used deterministic (time-domain) DCM for resting-state fMRI^39^. Our design required MRS-derived metabolite concentrations (Glu, Glx, GABA, and their ratios) to enter the model as a driving input shaping the integrated neuronal and haemodynamic response over time. This is consistent with the classic time-domain, input-driven DCM framework^39^.

#### Region of interest selection and times series extraction

Four ROIs were defined using anatomical masks in MNI space: the left lateral parietal (LPL), the medial prefrontal cortex (mPFC), the posterior cingulate cortex (PCC), and the right lateral parietal (RPL) (Supplemental Table 2). These regions were selected on the basis of their well-established roles as core DMN hubs and their consistent involvement in resting-state dysconnectivity across psychosis-spectrum conditions^18,20^. Regional time series (first eigenvariate, adjusted for effects of no interest) were extracted as VOIs; participants without activation in all four ROIs were excluded^30^. Please see the supplementary material for details.

#### Model space definition

Following Dauvermann et al. (2013)^40^, the intrinsic connectivity structure (Matrix A) was specified as a fully connected network across all four DMN nodes, consistent with established resting-state findings^18,20^, and Matrices B and C regressors were scaled by individual neurometabolite values. We constructed 32 models spanning three families: Baseline (one- vs two-state; Figure 1A), Direct Input (neurometabolite type — Glutamate, Glx, GABA, Glutamate/GABA, Glx/GABA — into mPFC or PCC; Figure 1B), and Modulatory (Figure 1C). Please see supplementary material for detailed justification.

**Figure 1.**
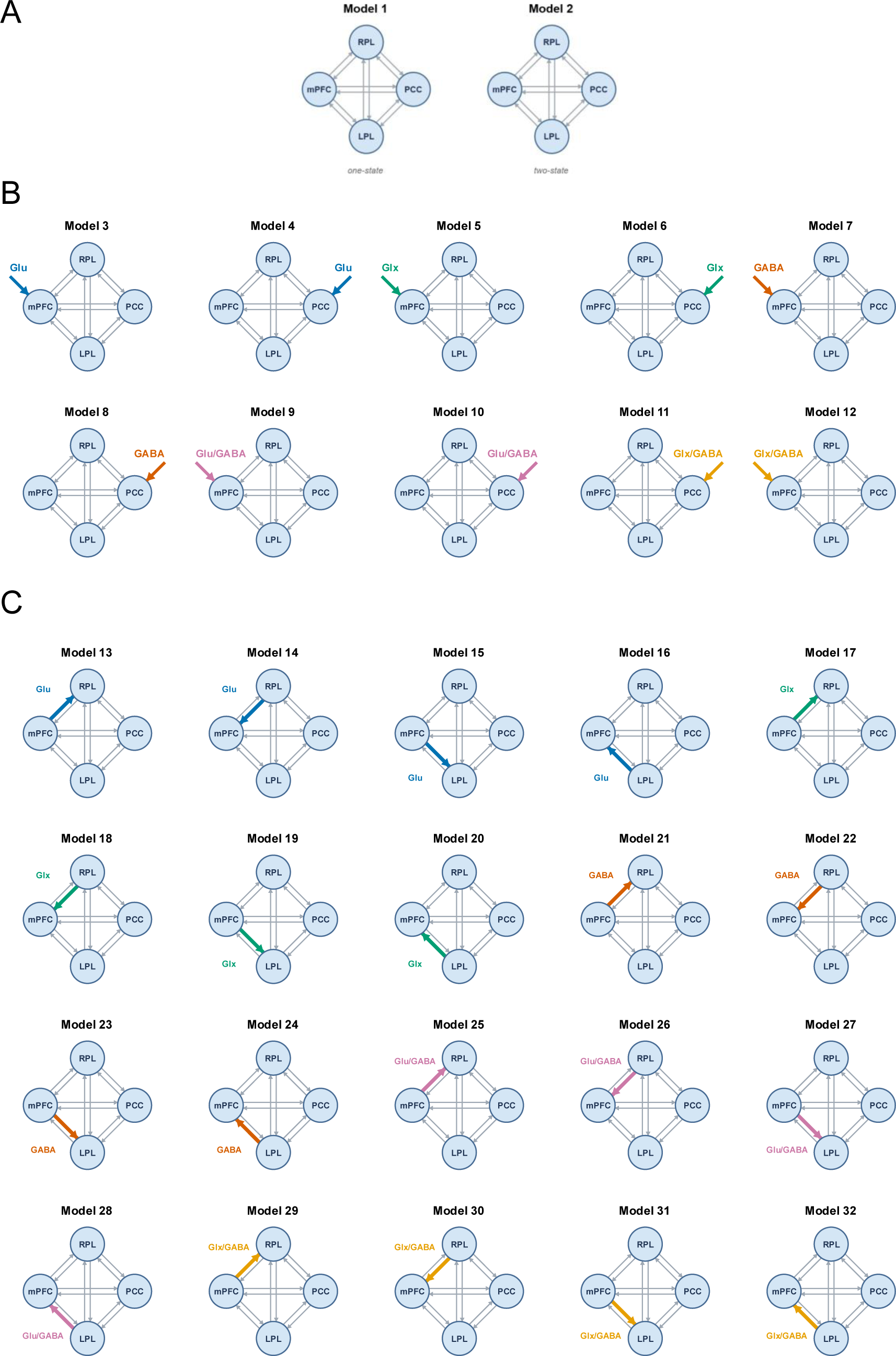
Model space. A. Baseline family: one-state model (model 1) versus two-state model (model 2). Models 1 and 2 served as baseline references: Model 1 used a one-state neural model with no driving input (C = 0) or modulatory connection (B = 0). Model 2 used a two-state neural model with no driving input or modulatory connection, enabling a direct comparison of state-model complexity in the absence of neurometabolite constraints B. Direct Input family: direct input region (mPFC or PCC) and neurometabolite type (Glutamate, Glx, GABA, Glutamate/GABA, Glx/GABA); (models 3 – 12). Models 3–12 tested direct driving input (C matrix) into either the mPFC or the PCC, scaled by one of five neurometabolite measures: Glutamate (Glu; models 3–4), Glutamate+Glutamine (Glx; models 5–6), GABA (models 7–8), the Glu/GABA ratio (models 9–10), or the Glx/GABA ratio (models 11–12). All models in this family used two-state dynamics. C. Modulatory family: neurometabolite type (Glutamate, Glx, GABA, Glutamate/GABA, Glx/GABA) as modulatory inputs (models 13 – 32). Models 13–32 tested the modulatory connection in the B matrix, testing four candidate modulated connections: RPL→mPFC, mPFC→RPL, mPFC→LPL, and LPL→mPFC, crossed with the five neurometabolite types (Glu, Glx, GABA, Glu/GABA, Glx/GABA) (Figure 1C). All models in this family also used two-state dynamics. Abbreviations. GABA, γ-Aminobutyric acid; Glu, Glutamate; Glx, Glutamate+Glutamine; mPFC, LPL, left parietal lobe; medial prefrontal cortex; PCC, posterior cingulate cortex; RPL, right parietal lobe.

### Bayesian Model Selection – Model-level and Family-level

To identify the optimal model at the group level, we performed random-effects Bayesian Model Selection (BMS) across all participants and all 32 models. BMS was conducted at both the individual model level and the family level. Family-level BMS was performed under the random-effects framework using Bayesian Model Averaging across the winning family. The key output metrics were exceedance probability (the probability that a given model or family is more likely than any other, given the group data) and posterior model probability. An exceedance probability exceeding 0.95 was used as the threshold for strong model evidence. The winning model identified by BMS was retained for all subsequent group-level analyses.

### Post-hoc Data Analysis – Relationships between DCM Parameters and Subclinical Symptoms

#### Parametric Empirical Bayes analysis

To examine between-participant sources of variability in effective connectivity parameters, parametric empirical Bayes (PEB) analysis was conducted on the winning model across all participants with complete data. PEB provides a hierarchical Bayesian framework in which second-level covariates are modelled as systematic influences on first-level DCM parameters, accounting for uncertainty in individual participant estimates through their posterior covariance matrices^39^. Please see supplementary material for details, including for the following Bayesian model reduction (BMR) and Bayesian model average (BMA) steps.

#### Correlation analysis between DCM parameters and subclinical measures

To complement the PEB analysis, Spearman rank correlations^41^ (robust to outliers/non-normality) were computed for the optimal model between the six A-matrix connections for the region receiving the driving input and the three subclinical symptom scores (21 correlations total; seven parameters × three measures, using raw scores). Bonferroni correction^42^ was applied (α = 0.05/21 ≈ 0.0024); uncorrected associations at p < 0.05 were reported as exploratory.

## Code availability

All code is publicly available on GitHub at: https://github.com/asadpouretal/two-state-dcm-ei-balance.

## Results

### Demographic and subclinical data

Demographic and subclinical data are presented in Table 1. No significant differences between females and males were observed for any of the demographic or subclinical data.

**Table 1.** Demographic and subclinical symptom details.

| | Females | Males | $t/\chi^2$ | P-value |
| --- | --- | --- | --- | --- |
| N (%) | 12 (50%) | 12 (50%) | - | - |
| Age; mean years (SD) | 23.08 (3.29) | 22.46 (4.96) | $t = -0.364$ | 0.719 |
| IQ; mean (SD) | 102.45 (6.98) | 98.27 (11.20) | $t = -1.051$ | 0.306 |
| Handedness N (right : left) | 11:1 | 11:1 | $\chi^2 = 2.182$ | 0.140 |
| SIPS; severity of psychotic experiences; mean (SD) | 1.25 (1.29) | 1.75 (1.48) | $t = 0.881$ | 0.388 |
| HAM-D; severity of depressive symptoms; mean (SD) | 3.5 (5.23) | 4.92 (3.60) | $t = 0.773$ | 0.448 |
| STAI; severity of anxiety symptoms; mean (SD) | 36.79 (8.96) | 35.25 (9.67) | $t = -0.405$ | 0.689 |

Abbreviations. HAM-D, Hamilton Depression Rating Scale; SIPS, Structured Interview for Psychosis-Risk Syndromes; SD, standard deviation; STAI, State-Trait Anxiety Inventory.

### Neuroimaging data

#### Magnetic Resonance Spectroscopy

Males showed significantly higher levels of Glutamate (females: mean = 18.07, SD = 1.37; males: mean = 20.00, SD = 1.03; *t* = 3.620, *p* < 0.001), GABA (females: mean = 3.42, SD = 0.18; males: mean = 3.54, SD = 0.12; *t* = 2.135, *p* = 0.044), and Glutamate/GABA (females: mean = 5.30, SD = 0.43; males: mean = 5.60, SD = 0.30; *t* = 2.018, *p* = 0.055) compared to females (Table 2). No further significant group differences were found. Independent-samples t-tests showed that males had a significantly higher white matter (WM) fraction than females (t(23) = −2.29, p = .031), whereas grey matter (GM) (p = .774) and cerebrospinal fluid (CSF) (p = .076–.079) did not differ significantly by sex. After controlling for WM, sex differences in Glutamate remained significant (F(1,22) = 18.30, p < 0.001, R² = .45), with WM itself not reaching significance (F(1,22) = 3.68, p = 0.068). The sex difference in GABA was no longer significant once WM was covaried (F(1,22) = 3.49, p = .075), whereas the sex difference in the Glutamate/GABA ratio became significant after adjustment (F(1,22) = 7.18, p = .014, R² = .26), with WM again non-significant (F(1,22) = 3.22, p = .087).

**Table 2.** Sex differences - Neurometabolite levels in the dorsal anterior cingulate cortex.

| | Females<br>(n = 12) | Males<br>(n = 12) | $t/U$ | P-value |
| --- | --- | --- | --- | --- |
| Glutamate, mean (SD) | 18.07 (1.37) | 20.00 (1.03) | $t = 3.620$ | $< 0.001^{**}$ |
| Glx, mean (SD) | 7.84 (0.82) | 8.34 (0.56) | $t = 1.828$ | 0.081 |
| GABA, mean (SD) | 3.42 (0.18) | 3.54 (0.12) | $t = 2.135$ | 0.044* |
| Glutamate/GABA, mean (SD) | 5.30 (0.43) | 5.60 (0.30) | $t = 2.018$ | 0.055 |
| Glx/GABA, mean rank | 10.58 | 14.42 | $U = 49.00$ | 0.184 |

Abbreviations. GABA, γ-Aminobutyric acid; Glx, Glutamate+Glutamine; mPFC, LPL, left parietal lobe; medial prefrontal cortex; PCC, posterior cingulate cortex; RPL, right parietal lobe.

### Bayesian Model Selection

#### Model-level results

Random-effects BMS was performed across the full 32-model space using data from all the participants. Model 11 – a two-state DCM with a fully connected intrinsic (A) matrix, no modulatory (B) connections, and direct (C) input of the Glx/GABA ratio into the PCC – emerged as the clear winning model (Figure 2A). Model 11 achieved an exceedance probability (XP) of 0.96 (96.0%), exceeding the conventional 0.95 threshold for strong evidence, and accounted for the largest share of the posterior model probability (PP = 0.79; 78.9%). The XP of the second-ranked model, Model 12 (Glx/GABA ratio input to the medial prefrontal cortex, MPFC), was more than 25-fold lower (XP = 3.6%) (Table 3A).

**Figure 2.**
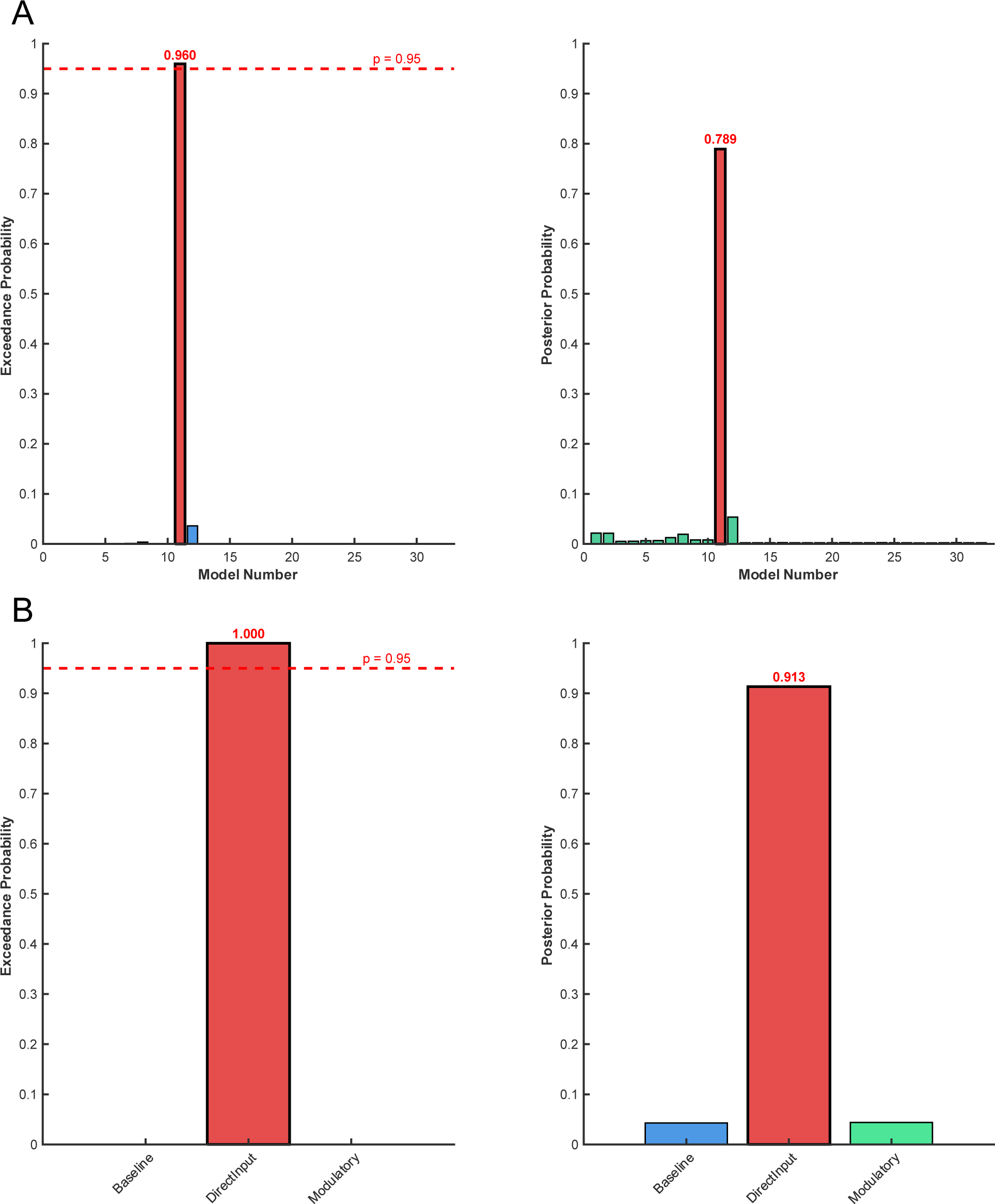
Bayesian Model Selection results. **A. Model-level random-effects Bayesian Model Selection across the 32-model space (N = 20).** Left: exceedance probabilities for each candidate model; Model 11 (red bar) was the only model to exceed the 0.95 threshold (dashed red line), with an exceedance probability of 0.960. Right: posterior probabilities for each model; Model 11 again accounted for the greatest share of the evidence (0.789). **B) Family-level random-effects Bayesian Model Selection (N = 20).** Left: exceedance probabilities for the Baseline (Models 1–2), DirectInput (Models 3–12), and Modulatory (Models 13–32) families; the DirectInput family (red bar) achieved a perfect exceedance probability of 1.000, well above the 0.95 threshold (dashed red line). Right: posterior probabilities for each family, confirming the dominance of the DirectInput family (0.913).

**Table 3.**
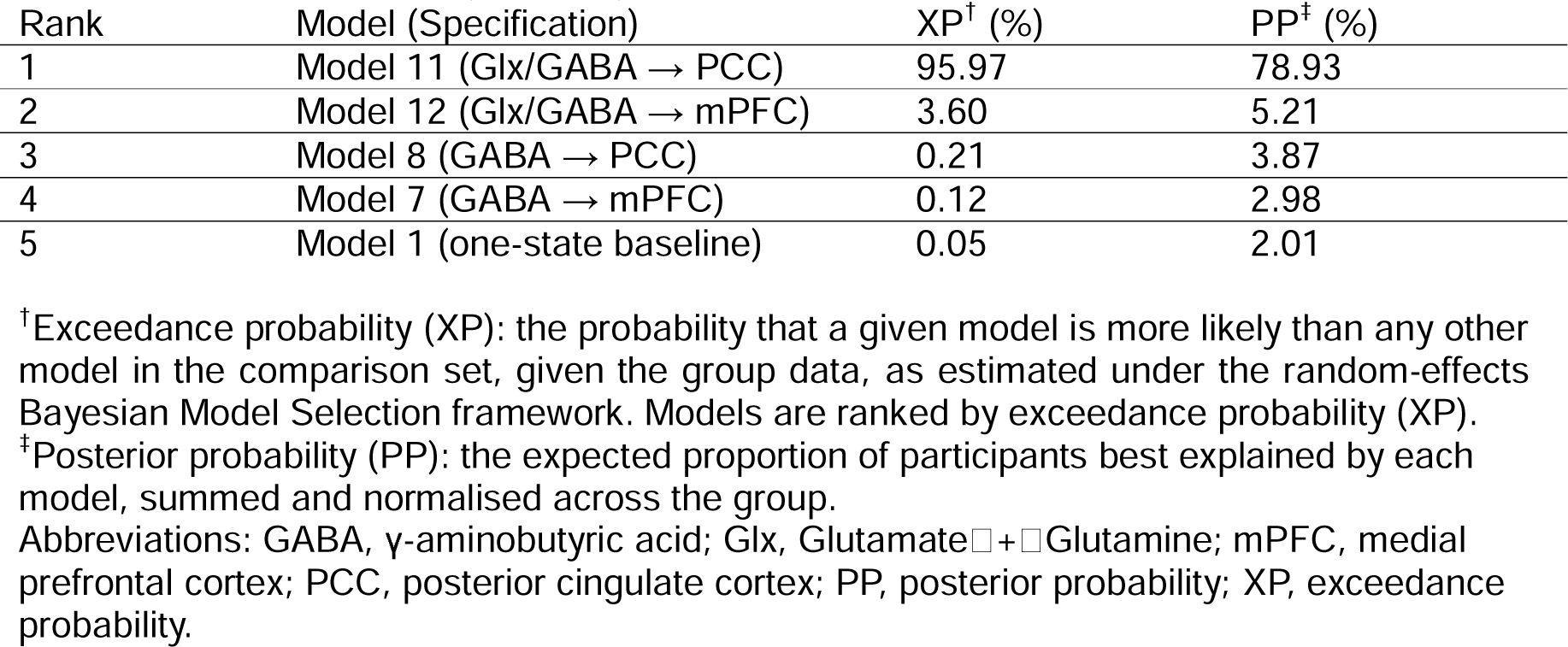
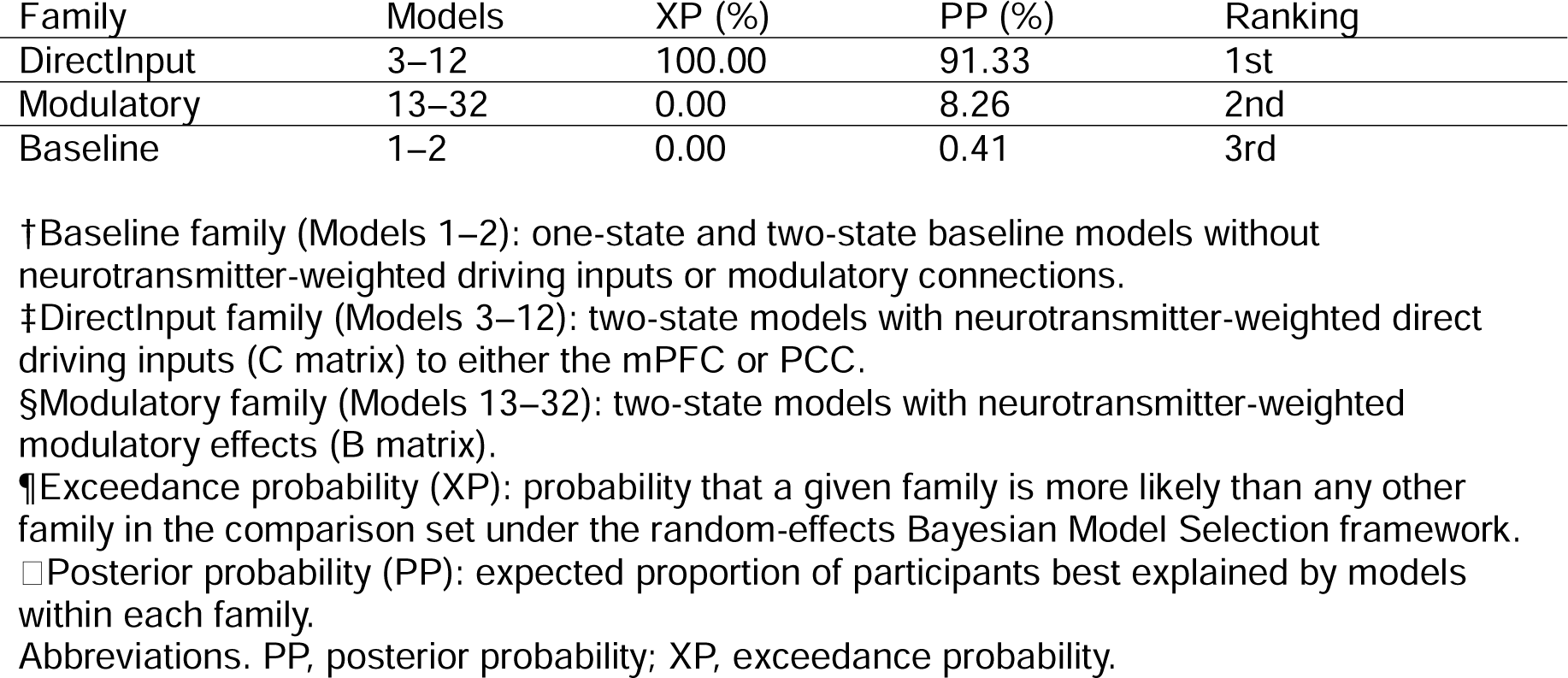
Model-level and family-level Bayesian Model Selection results (N = 20). A. Top five ranked models from random-effects Bayesian Model Selection across the 32-model candidate space (N = 20). B. Family-level random-effects Bayesian Model Selection across the three predefined model families (N = 20).

Across the top five ranked models, three patterns were apparent. First, the three best-performing models (Models 11, 12 and 8) all used a neurometabolite ratio (Glx/GABA) or an absolute neurotransmitter concentration (GABA) as the driving input, rather than no input at all or a modulatory connection. Second, the PCC was favoured over the mPFC as the target of direct driving input among the top-ranked models. Third, every model among the top five used two-state, rather than one-state, neuronal dynamics, with the one-state baseline model (Model 1) ranking only fifth and accounting for a negligible proportion of the evidence (XP = 0.05%; PP = 2.0%). No model with modulatory connection was among the five most likely models.

#### Family-level results

To test whether neurometabolite information was better explained as a direct driving input, as a modulator of existing connections, or was unnecessary altogether, the 32 models were partitioned into three *a priori* families – Baseline (Models 1–2), Direct Input (Models 3–12), and Modulatory (Models 13–32) – and family-level random-effects BMS was performed (Figure 2B). The Direct Input family was identified as the winning family with an exceedance probability of 1.00 (100%) and a posterior probability of 0.91 (91.3%), providing decisive evidence over both the Modulatory family (XP = 0.00%; PP = 8.3%) and the Baseline family (XP = 0.00%; PP = 0.4%) (Table 3B). This family-level result converges with the model-level finding above: neurometabolite ratios and concentrations appear to act predominantly as direct driving inputs into DMN regions (C matrix), rather than as modulators of the strength of existing intrinsic connections (B matrix).

### Parametric Empirical Bayes Analysis

Of the 20 participants with a valid winning-model (Model 11), one participant had incomplete subclinical data and was excluded from all second-level analyses, yielding a final sample of N = 19 (11 male, 8 female) for the PEB and correlation analyses.

#### Full model: Biological sex and subclinical symptom severity

A second-level PEB model was estimated on the A-matrix (intrinsic connectivity) and C-matrix (driving input) parameters of Model 11. Following model reduction, no DCM parameter showed strong evidence (posterior probability, Pp > 0.95) for an effect of biological sex, severity of psychotic experiences, or severity of subclinical levels of depression (Supplemental Figure 2). For severity of subclinical levels of anxiety, several A-matrix connections showed posterior probabilities of approximately 0.50, but the corresponding posterior effect sizes were negligible in magnitude and did not approach conventional thresholds for evidence. Overall, there was no clear evidence that current subclinical symptom severity robustly modulated the effective connectivity parameters of the winning DCM.

#### Post-hoc model: Biological sex only

Given the *a priori* interest in biological sex differences, a post-hoc PEB analysis was conducted. This analysis did not identify evidence for an effect of biological sex on effective connectivity: no parameter reached a liberal posterior probability threshold of 0.49, and BMR did not retain a biological sex-related effect for any connection. This null result is consistent with the absence of a biological sex effect observed in the full PEB model reported above.

Taken together, the PEB and correlation analyses (refer to supplementary materials for correlation analysis results) converge on a consistent pattern: within this proof-of-concept sample, biological sex, and severity of psychotic experiences and subclinical levels of depression and anxiety) were not robustly associated with individual differences in effective connectivity of the winning DCM (Model 11).

## Conclusion

This proof-of-concept study examined whether excitatory/inhibitory balance in the regions of interest underlies resting-state DMN effective connectivity in healthy young adults. A two-state model with the Glx/GABA ratio as direct input into the PCC provided the best fit, and higher dACC Glx/GABA ratios were associated with greater severity of psychotic experiences — supporting the hypothesis that excitatory/inhibitory balance underlies stable resting-state effective connectivity^27,28^ and offering a mechanistic account of how this balance may translate into network-level dysfunction.

The main finding is that the two-state DCM model with the Glx/GABA ratio as a direct input into the PCC provided the best explanation of resting-state effective connectivity within the DMN, outperforming both the one-state model and models using alternative metabolite inputs. This has two key implications: it demonstrates that neurochemical excitatory/inhibitory balance is not merely a local property of individual cortical regions but is functionally integrated with large-scale network connectivity, as previously theorised in computational studies of neurotransmitter homeostasis^27,28^; and it identifies the PCC — a key DMN hub with an established role in default-mode suppression^43^ — as the region through which Glx/GABA exerts its influence on the DMN, which is neurobiologically plausible^44,45^ given its consistent implication in altered resting-state connectivity across schizophrenia-spectrum conditions^46^.

The two-state DCM formulation’s explicit separation of excitatory and inhibitory neuronal populations provides a biophysically interpretable framework for investigating excitatory/inhibitory balance at the network level, unlike standard one-state models. That the three best-fitting models all incorporated GABA or its ratios further validates this approach, aligning with computational modelling work showing that local cortical excitatory/inhibitory homeostasis sustains healthy brain network function^27–29^, and extending this evidence to a non-invasive, in-vivo human context.

Using the Glx/GABA ratio (capturing Glutamate and its precursor Glutamine alongside GABA) rather than Glutamate/GABA alone reflects evidence that Glutamine contributes meaningfully to excitatory tone via the Glutamate/Glutamine cycle^47,48^, and echoes prior associations between Glx/GABA and DMN dysconnectivity in healthy participants^26,49^ and high-schizotypy individuals^50^, which lends further validity to our network-level excitatory/inhibitory measure.

Higher dACC Glx/GABA ratios were related (trend-level) to greater severity of psychotic experiences, suggesting both excitatory and inhibitory levels are relevant to subclinical symptom expression, consistent with the broader excitatory/inhibitory balance literature at the high-risk stage of psychosis^7,50^, and potentially indicating a neurochemical pattern already present at the subclinical/risk stage. These findings are exploratory and correlational; caution is warranted in drawing causal inferences.

Males showed significantly higher dACC Glutamate, GABA, and Glutamate/GABA ratio than females, though exploratory sex-difference analyses of excitatory/inhibitory balance and effective connectivity were non-significant. Given known sex differences in prefrontal cortical excitability^51^ and inhibition^52^, future adequately powered studies should systematically examine sex as a moderator of the excitatory/inhibitory balance–DMN connectivity relationship; the present analyses were exploratory and underpowered for sex-specific effects. The robustness of the Glutamate sex difference after WM adjustment suggests this effect is not simply a by-product of the underlying sex difference in tissue composition. In contrast, the GABA sex difference dropped below threshold once WM was accounted for, raising the possibility that it was at least partly related to the tissue-composition difference between sexes rather than reflecting an independent effect on GABA itself. The Glutamate/GABA ratio showed the opposite pattern, becoming more significant after adjustment, which is consistent with WM absorbing some of the unexplained variance in the ratio without itself being a significant predictor, sharpening rather than accounting for the sex effect.

Several methodological strengths merit acknowledgement. First, integrating MRS-derived Glx/GABA ratios directly into the DCM model as empirical priors — rather than correlating MRS and connectivity measures post hoc — is a novel, biophysically principled approach that improves biological interpretability and mechanistic precision; to our knowledge, this is among the first applications of this approach in healthy humans with psychotic experiences. Second, two-state DCM’s explicit separation of excitatory and inhibitory populations, combined with evidence-based BMS model selection, is better suited to testing excitatory/inhibitory hypotheses than conventional one-state models. Third, our dimensional, continuum-based sampling of subclinical psychotic, anxious, and depressive experiences (consistent with RDoC^53^) avoids confounds associated with clinical samples (e.g., medication, illness chronicity, diagnostic heterogeneity). Fourth, the multimodal, single-session design with quality control procedures, and well-justified dACC voxel placement, provides methodological robustness. Finally, comparing Glutamate/GABA against Glx/GABA in the model space identified Glx/GABA as the more informative predictor, adding mechanistic granularity regarding Glutamate and Glutamine’s relative contributions.

Several limitations must be acknowledged. First, the small sample limits statistical power and generalisability, precludes robust multiple-comparison correction, and means associations should be treated as hypothesis-generating; effect size estimates are nonetheless informative for designing adequately powered future studies. Second, the cross-sectional design precludes causal inference about the direction (or bidirectionality) of the dACC excitatory/inhibitory balance– DMN connectivity–symptom relationship; longitudinal designs with repeated assessments, e.g. before/after an experimental stress perturbation, are needed. Third, the sample was sex-balanced but not powered for sex-specific effects. Fourth, while biophysically motivated, two-state DCM relies on simplifying assumptions linking neural population connectivity to the BOLD signal; its haemodynamic observation model is less neurophysiologically detailed than DCM for EEG/MEG, so parameter estimates should be interpreted cautiously, and future studies combining fMRI-based DCM with electrophysiological measures may add complementary insight. Fifth, Glutamate/GABA concentrations are a static, voxel-pooled proxy for excitatory/inhibitory balance and cannot be directly compared to circuit-level synaptic release, receptor occupancy, or firing rates, nor do they capture receptor subtype or density (e.g., NMDA receptors, parvalbumin interneurons) — limitations we sought to partly address through improved Glx/GABA reliability via spectral editing, optimised tissue-composition modelling, and integration with two-state DCM as an inferential measure.

### Clinical Implications

Despite its preliminary nature, this study has potential clinical implications. That a network-level excitatory/inhibitory measure combining dACC Glx/GABA with PCC-centred connectivity was sensitive to subclinical psychotic experiences suggests utility as an early psychosis vulnerability marker, potentially complementing existing risk stratification approaches, which have been hampered by at-risk-state heterogeneity and a lack of neurobiologically grounded, dimensional measures^54^. As excitatory/inhibitory balance is modifiable — pharmacologically (e.g., NMDA receptor modulation) and psychologically (e.g., stress-regulation interventions) — this network-level measure may also serve as a target engagement biomarker in future intervention studies^55^.

In conclusion, excitatory/inhibitory balance in the PCC — defined by the dACC Glx/GABA ratio as an empirical prior in a two-state DCM — predicted resting-state DMN effective connectivity, and higher Glx/GABA ratios were associated with greater psychotic experience severity. These proof-of-concept findings support a mechanistic pathway linking neurochemical excitatory/inhibitory balance to network-level dysfunction and subclinical symptom expression, consistent with a psychosis-continuum framework. Replication in larger, adequately powered samples with experimental excitatory/inhibitory manipulation is a key next step.

## Author contributions

Conceptualisation: A.A., M.R.D. Methodology: A.A., M.W., M.R.D. Formal analysis: A.A., M.W., M.R.D. Investigation: A.M., J.C., A.G., P.B., I.M. Writing – original draft: A.A., M.R.D. Writing – review & editing: A.M., J.C., A.G., P.B., I.M., M.W. Supervision: M.R.D. Funding acquisition: M.R.D.

## Acknowledgments

This work was supported by the University of Birmingham as a start-up package to M.R.D. The authors thank the MRI radiographer and technicians at the Centre for Human Brain Health for assistance with data acquisition. We are grateful to all participants who took part in this study.

