## Supplementary Material for "Two-State Effective Connectivity Predicts the Effect of Anterior Cingulate Cortex Excitatory/Inhibitory Balance on Posterior Cingulate Cortex"

Short title: Two-State Modelling of Effect of ACC Excitatory/Inhibitory Balance on PCC

Abdoreza Asadpour[1], Aanya Malaviya[2], Alan George[2], Piyali Bhattacharya[2], Ifigeneia Manitsa[2], Martin Wilson[3], Maria R Dauvermann[2]

Institutional affiliations:

[1]Sussex Neuroscience, School of Life Sciences, University of Sussex, Brighton, UK

[2] Institute for Mental Health, School of Psychology, University of Birmingham, Birmingham, UK

[3] Centre for Human Brain Health, School of Psychology, University of Birmingham, Birmingham, UK

### Methods

#### Participants

Thirty-seven young healthy adults participated in this study. Participants were recruited via the University of Birmingham, public places and the online research platform Call for Participants (https://www.callforparticipants.com). All participants were aged between 18 and 35 years of age, healthy (e.g., did not have a clinical diagnosis for healthy condition and did not receive treatment for a health condition, including a psychotic disorder, anxiety and depression). Exclusion criteria included presence of a documented history of neurological disorders (e.g., epilepsy), mental health conditions (including substance misuse in the last six months), an estimated intelligence quotient (IQ) less than 70, a lifetime history of head injury causing loss of consciousness for more than five minutes, reported pregnancy or lactation, and any contra-indications for MRI scanning (e.g., mental implants or claustrophobia). All participants were balanced by sex (19 females and 18 males). Seventeen participants were excluded from this analysis due to missing data or failed data quality control on subclinical data (n = 1), no neuroimaging data (n = 8), MRS spectra (n = 1), failed volume of interest (VOI) extraction (n = 7). All participants provided written informed consent in accordance with the guidelines of the local Ethics Committee at the University of Birmingham.

#### Data Collection

##### Demographic data

Typical demographic details have been collected. The Wechsler Abbreviated Scale of Intelligence – Second Edition[1]. The following four subtests were administered: Vocabulary, Similarities, Block Design and Matrix Reasoning. This is a brief reliable IQ estimate that was normed on a nationally representative sample. It provides a quick, reliable and valid estimate of IQ when administration of a full battery is not feasible or necessary; particularly useful for research applications; easy to learn and administer. This edition shows high reliability across adult research samples with test-retest reliability coefficients between 0.80 and 0.90[2].

##### Subclinical symptom assessment

Psychotic experiences and subclinical affective symptoms were assessed by trained researchers using established, interview-based clinical instruments. The severity of psychotic experiences was evaluated using the Structured Interview for Psychosis-Risk Syndromes (SIPS)[3], which provided scores across four subscales: positive symptoms, negative symptoms, disorganised symptoms, and general symptoms. The SIPS demonstrates excellent inter-rater reliability, predictive validity, and specificity in identifying individuals at clinical high risk for psychosis[4]. Subclinical levels of depression were measured using the Hamilton Depression Rating Scale[5], an instrument widely used to quantify the severity of depressive symptoms across affective, somatic, and cognitive domains. This scale demonstrates robust test-retest and inter-rater reliability coefficients[6]. State and trait dimensions of subclinical levels of anxiety were assessed using the State-Trait Anxiety Inventory (STAI)[7], which distinguishes between anxiety as a transient emotional state and anxiety as a stable dispositional characteristic. All interviews were conducted by researchers trained to criterion on each instrument prior to data collection, and interrater reliability was established before the study commenced. The STAI has excellent psychometric properties, consistently demonstrating high levels of internal consistency and reliability across various demographics, cultural groups, and translated versions[8].

#### Neuroimaging data acquisition

All participants underwent a single neuroimaging session. Brain imaging was carried out on a 3 Tesla Siemens Magnetom Prisma (Siemens Healthcare, Erlangen, Germany) system using a 32-channel receiver head coil array at the Centre for Human Brain Health, University of Birmingham. The brain imaging data were acquired in the following order: T1-weighted scan, rs-fMRI scan, single-voxel MRS (semi-LASER and MPRESS sequences).

##### Structural Magnetic Resonance Imaging

A T1-weighted MRI scan was acquired sagittally with a 3D-MPRAGE sequence: FOV = 208 x 256 x 256 mm, resolution = 1 x 1 x 1 mm, TE / TR = 2ms / 2000 ms, inversion time = 880 ms, flip angle = 8˚, and GRAPPA acceleration factor  = 2 (4 min 54 s scan duration).

##### Magnetic Resonance Spectroscopy

Single-voxel MRS was acquired in the dACC using semi-LASER[9] (Glutamate; 20 x 25 x 20 mm (RL x AP x FH) voxel) and MEGA-PRESS[10] (GABA; 30 x 35 x 25 mm (RL x AP x FH) voxel) sequences, with standardised voxel placement in the dACC.

One hundred and ninety-two transients were acquired with a 90˚ flip angle and an 8-step phase cycling scheme; TR / TE = 2000 / 28 ms and VAPOR water suppression (6 min 38 s scan duration). Four transients were also acquired without VAPOR water suppression. $B_0$ shimming was performed using the ‘brain’ method implemented by the vendor.

The GABA-edited MEGA-PRESS sequence (MPRESS) with TR / TE = 2000 / 68 ms and 128 averages (8 min 56 s scan duration) was used. Editing pulses of 55 Hz bandwidth were applied alternately at 1.90 ppm (ON condition) and 7.50 ppm (OFF condition). Subtraction of the ON sub-spectrum from the OFF sub-spectrum generates a difference spectrum in which the GABA resonance at 3.0 ppm is revealed. $B_0$ shimming was performed using the ‘brain’ shim method implemented by the vendor. An additional water-reference scan (1 average, VAPOR suppression disabled) was acquired immediately after the MPRESS sequence with identical voxel placement and sequence parameters (TE = 68 ms) to enable absolute metabolite quantification. As the GABA editing pulse at 1.9 ppm also co-edits a macromolecule resonance at 3.0 ppm, the resulting GABA signal reflects GABA plus macromolecules (GABA+ [11]).

Given the well-established poor agreement between Glx and Glutamate measured from MEGA-PRESS OFF sub-spectra and difference spectra relative to dedicated short-echo PRESS acquisitions[12], Glutamate was quantified from the semi-LASER sequence, whereas Glx was quantified from the MPRESS sequence. All MRS data from both the semi-LASER and the MPRESS sequences were exported and converted to NifTI MRS format[13] as 2048 complex data points were acquired with a spectral width of 2000 Hz.

Voxel placement followed a standardised protocol. Cortical landmarks were used to identify the coronal slice and define the voxel centre, after which the voxel was rotated in the transverse and sagittal planes to achieve the final positioning. The placement of the dACC voxel is illustrated in supplemental Figure 1, with representative spectra for GABA and Glutamate levels.

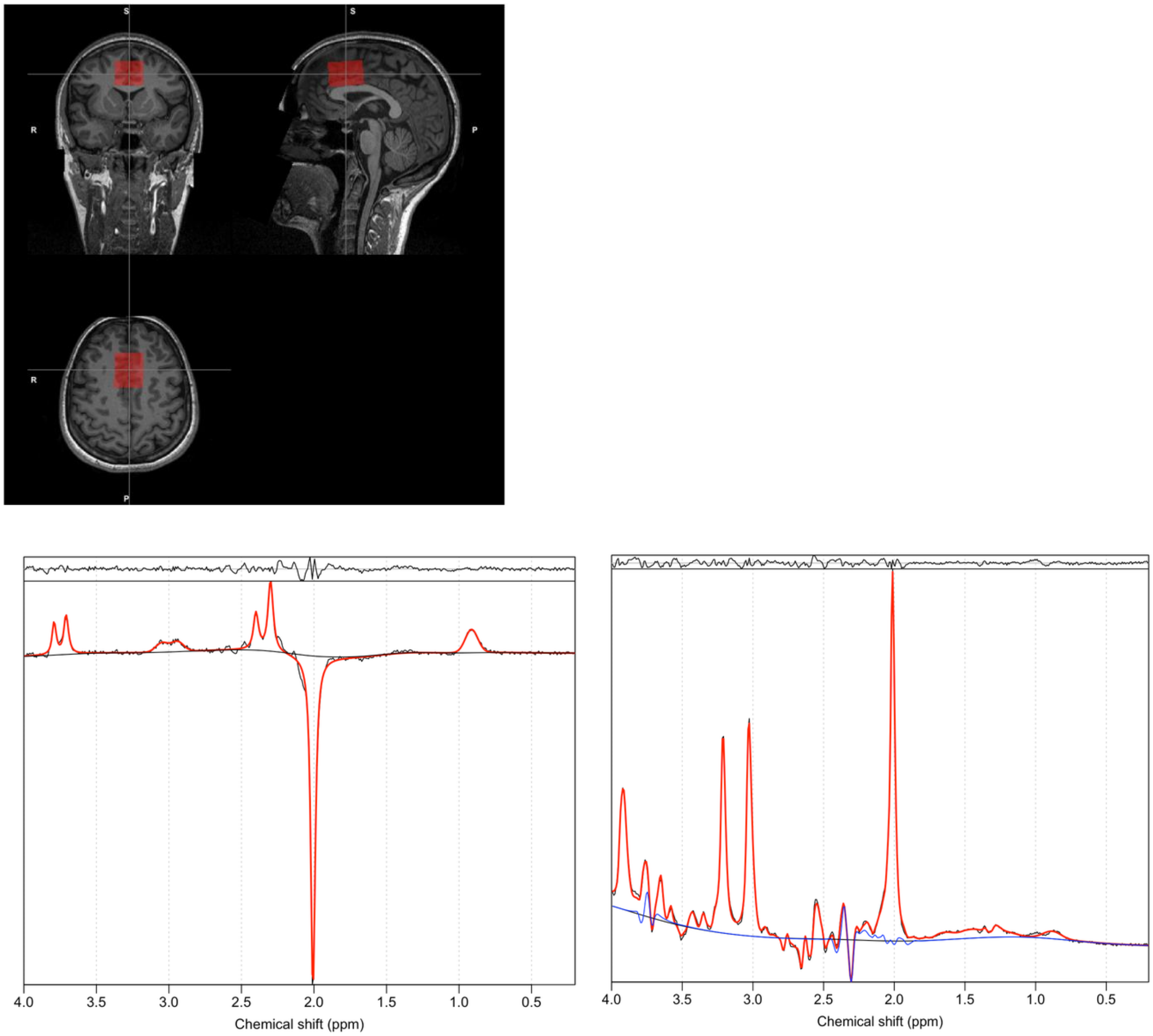


**Supplemental Figure 1. Voxel placement and representatively fitted MRS spectra for GABA and Glutamate levels in the dorsal anterior cingulate cortex.**
**A.** Placement of MRS voxel in the dorsal anterior cingulate cortex and $^{1}$H-MRS spectrum fitted by Spant for:
**B**. GABA
**C**. Glutamate based on edited MEGA-PRESS sequence.

#### Resting-state functional Magnetic Resonance Imaging

Rs-fMRI data was acquired using a multi-band accelerated GRE-EPI sequence with an isotropic voxel resolution of 2.5 mm over a 210 x 210 x 142.5 mm FOV (AP x LR x FH). 394 fMRI volumes (each containing 57 transverse slices) were acquired with a TR = 1500 ms, TE = 35 ms; 71° excitation flip angle; A>>P phase encoding direction; and multi-band acceleration factor = 3 (10 min 1 s scan duration).

### Neuroimaging data analysis

#### Magnetic Resonance Spectroscopy Data Analysis

Both the semi-LASER and MPRESS MRS data were corrected for single-shot phase and frequency instability using the RATS method[9] as a preprocessing step. Spectral fitting was performed with ABfit-reg[10] using a simulated basis-set matched to the MRS acquisition parameters. For MPRESS, both the edited and edit-off data were analysed, allowing estimates of GABA+ alongside metabolite levels estimated from the conventional PRESS data (edit-off scans). The proportion of white matter, grey matter and CSF present in each MRS acquisition volume was calculated from the T1 anatomical data following segmentation with FSL FAST[14] (please see results for these tissue proportions in Supplemental Table 1). Metabolite levels were scaled relative to water reference scans, incorporating the segmented proportions of grey/white-matter and CSF, according to standard relaxation assumptions[15]. All MRS processing steps were implemented in the spant analysis package[16] developed for the R programming language.

**Supplemental Table 1. Tissue proportion results for the dorsal anterior cingulate cortex voxel (N = 24)**

| | % Mean | % SD |
|---|---|---|
| WM | 36.30 | 3.70 |
| GM | 48.96 | 3.27 |
| CSF | 14.84 | 3.76 |
| Other | 0 | 0 |

Abbreviations. CSF, cerebrospinal fluid; GM, grey matter; WM, white matter.

### Resting-state functional Magnetic Resonance Imaging Data Analysis

#### Pre-processing of rs-fMRI data

Pre-processing and first-level general linear model of the rs-fMRI data was performed using Statistical Parametric Mapping (SPM12; Wellcome Trust Centre for Neuroimaging, London, UK; https://www.fil.ion.ucl.ac.uk/spm) in MATLAB (R2022a; The MathWorks Inc., Natick, MA).

Slice timing correction was applied to all functional volumes, co-registered to native space of the structural data, before the volumes were realigned, segmented, normalised to MNI space, and spatially smoothed with an isotropic 8 mm full-width at half-maximum (FWHM) Gaussian kernel to compensate for residual variability in functional anatomy after spatial normalisation and to facilitate the application of Gaussian random field theory for adjusted statistical inference.

A first-level general linear model (GLM) was applied using 36 confound regressors in line with established guidelines[17,18]. The 36 regressors comprised six motion parameters, three tissue-based signals (white matter, cerebrospinal fluid, and global signal), and their respective first-order temporal derivatives and quadratic expansions. Global signal regression was included to minimise the contribution of widespread noise to regional time series estimates. Low-frequency drift was removed by incorporating three discrete cosine basis functions into the GLM. Outlier time points were identified and modelled using stick regressors, applied at thresholds of 0.5 mm for frame displacement and 1.5 for standardised DVARS.

#### Two-state Dynamic Causal Modelling for fMRI

##### Background to DCM, two-state and stochastic DCM for fMRI

DCM is an established method for assessing inter-regional effective connectivity by modelling experimentally or spontaneously induced changes in neural activity[19]. In contrast to standard functional connectivity methods, which measure correlations between regional time series, DCM enables causal inference by modelling directed influences between brain regions at the neuronal level. It does so by estimating how the rate of change of neural activity in one region influences neural activity in other regions, using coupled differential equations that are linked to predicted BOLD responses via a biophysical haemodynamic forward model[19].

The standard bilinear formulation of DCM — referred to here as the one-state model — models each brain region as a single neuronal population, thereby conflating excitatory and inhibitory contributions to effective connectivity. The two-state DCM extends this framework by explicitly separating each region into excitatory and inhibitory neuronal subpopulations[20]. This biophysical distinction makes the two-state model better suited to testing hypotheses about excitatory/inhibitory balance, as it allows excitatory and inhibitory contributions to inter-regional connectivity to be estimated independently. This is particularly relevant in the context of the present study, in which we sought to examine how MRS-derived Glutamate/GABA ratios in the dACC modulate effective connectivity within the DMN.

In addition, we used spectral DCM for resting-state fMRI, which characterises the low-frequency (<0.1 Hz) cross-spectral density of spontaneous neuronal fluctuations by estimating the amplitude and spectral exponent of hidden neuronal states within each region of a predefined network[21].

Because spectral DCM uses a power-law form, it provides a deterministic and computationally efficient model that captures stochastic neural dynamics without requiring an explicit external input. Connectivity strengths are expressed in Hz, such that higher values correspond to faster induced responses in downstream regions. The resulting neuronal dynamics are transformed into predicted fMRI BOLD signals via a haemodynamic forward model[22].

#### Region of interest selection and times series extraction

Four ROIs were defined using anatomical masks in MNI space: the left lateral parietal (LPL), the medial prefrontal cortex (mPFC), the posterior cingulate cortex (PCC), and the right lateral parietal (RPL) (Supplemental Table 2). These regions were selected on the basis of their well-established roles as core DMN hubs and their consistent involvement in resting-state dysconnectivity across psychosis-spectrum conditions[23,24]. The regional time series were extracted as volumes of interest (VOIs) using the first eigenvariate of all voxels within each region, adjusted for effects of no interest. The condition of activation within each ROI had to be met for a subject to be included in the DCM analyses; subjects who did not show activation in all four ROIs satisfying the inclusion criteria were excluded from further analysis[19]. Haemodynamic response timing was aligned to the VOI acquisition time (T0 = RT × t0/t), and an echo time of 35 ms was used throughout, consistent with the rs-fMRI acquisition parameters described above.

**Supplemental Table 2.** Coordinates of the four default-mode network seeds

| Brain regions | Coordinates in MNI space | | |
|---|---|---|---|
| | x | y | z |
| mPFC | 1 | 55 | -3 |
| LPL, BA39 | -39 | -77 | 33 |
| RPL, BA39 | 47 | -67 | 29 |
| PCC | 1 | -61 | 38 |

Abbreviations. BA, Brodman area; LPL, Left Parietal Lobe; Medial PFC, medial Prefrontal Cortex; PCC, Posterior Cingulate Cortex; RLP, Right Parietal Lobe.

#### Model space definition

Following the approach of Dauvermann et al. 2013[25], who grounded their DCM model specification in neuroimaging evidence, deriving the intrinsic connectivity structure (Matrix A) from neuroimaging studies, and specifying the modulatory (Matrix B) and driving input (Matrix C) parameters from established studies, we adopted an analogous framework in the present study.

Specifically, the intrinsic connectivity structure (Matrix A) was specified as a fully connected network, allowing bidirectional influences between all four DMN nodes (LPL, mPFC, PCC, and RPL), consistent with established resting-state connectivity findings in healthy adults and individuals with psychotic disorders[23,24]. For modelling the driving input (Matrix C) and the modulatory connection (Matrix B), we drew on evidence from multimodal MRI studies in individuals with psychotic disorders that combined resting-state fMRI functional connectivity with MRS-derived Glutamate, Glx and GABA metabolites. Each of these studies used correlation analysis to relate resting-state functional connectivity findings with Glutamate or GABA concentrations. With respect to Glutamate and Glx, reduced Glutamate levels in the ACC have been associated with lower functional connectivity in the parietal lobe in first-episode psychosis (FEP)[26], while lower Glutamate levels in FEP were further linked to reduced within-region functional connectivity of the mPFC[27]. Glutamate concentrations in FEP showed a positive relationship with functional connectivity in the precuneus and a negative relationship with the medial frontal cortex[28]. Regarding GABA, lower GABA levels in the ACC were associated with altered functional connectivity in the parietal lobe in FEP[26], while an anti-correlation between GABA levels and within-region mPFC functional connectivity observed in healthy controls was absent in FEP[27]. GABA levels showed a positive relationship with IFG functional connectivity in FEP and with mPFC functional connectivity in healthy controls[28]. Furthermore, changes in GABA in the mPFC were correlated with alterations of DMN functional connectivity following eight weeks of treatment in FEP[29], and GABA levels in the dACC were negatively correlated with functional coupling within the DMN in healthy controls[30]. Finally, and most directly informing the present model, the Glx/GABA ratio has been associated with DMN functional connectivity in the parietal lobe and PCC in healthy controls[30], providing direct empirical support for the use of this ratio as a driver of PCC-centred DMN effective connectivity in the present study.

In summary, we constructed 32 models that addressed  state-model complexity (one-state model versus two-state models; Figure 1A), direct input region (mPFC or PCC; Figure 1B) neurometabolite type (Glu, Glx, GABA, Glu/GABA, Glx/GABA) (Figure 1B and Figure 1C), and modulatory connection (Figure 1C).

#### Post-hoc Data Analysis – Relationships between Magnetic Resonance Spectroscopy and Subclinical Symptoms

As a post-hoc exploratory analysis, we conducted a median split on Glx/GABA levels and examined bivariate correlations in the subgroup with low Glx/GABA ratios ($n = 12$). These results should be interpreted with caution given the small sample size and the exploratory, hypothesis-generating nature of this analysis. In this subgroup, Glx/GABA levels showed a trend-level positive correlation with severity of psychotic experiences (SIPS, positive symptoms; $r = 0.572$, $p = 0.052$, 95% CI [0.187, 0.823], bootstrap = 5,000), suggesting that higher Glx/GABA concentrations in the dACC were associated with greater psychotic symptom severity. No other significant correlations were found.

### Post-hoc Data Analysis – Relationships between DCM Parameters and Subclinical Symptoms

#### Parametric Empirical Bayes analysis

To examine between-participant sources of variability in effective connectivity parameters, parametric empirical Bayes (PEB) analysis was conducted on the winning model across all participants with complete data. PEB provides a hierarchical Bayesian framework in which second-level covariates are modelled as systematic influences on first-level DCM parameters, accounting for uncertainty in individual participant estimates through their posterior covariance matrices[31].

For PEB analysis, variables included biological sex (coded 0 = male, 1 = female), and continuous scores on three symptom measures: severity of psychotic experiences, severity of subclinical levels of depression, and severity of subclinical levels of anxiety. Continuous measures were mean-centred prior to entry into the design matrix.

The PEB design matrix comprised five columns: (1) a group mean (intercept), (2) sex, (3) severity of psychotic experiences (mean-centred), (4) severity of subclinical levels of depression (mean-centred), and (5) severity of subclinical levels of anxiety. The PEB model was estimated with *a priori* on the between-subject variability of all DCM parameters and estimating a full second-level covariance structure. The A matrix (intrinsic inter-regional connections) and C matrix (driving inputs from neurometabolite-scaled regressors) were jointly submitted to the PEB estimation.

Following PEB estimation, we applied Bayesian model reduction (BMR) which performs an exhaustive search over all combinations of second-level parameters (covariate–DCM parameter pairs), pruning those not supported by the data[32]. The resulting Bayesian model average (BMA) provides posterior probability estimates for each covariate effect on each DCM parameter. Effects were considered significant when their posterior probability exceeded 0.95[31]. To further examine sex-specific effects, a dedicated post-hoc PEB analysis was also conducted with biological sex as the sole covariate (in addition to the group mean), following the same estimation and model reduction procedure.

#### Correlation analysis between DCM parameters and subclinical measures

To complement the PEB analysis with a model-free characterisation of parameter–symptom relationships, Spearman rank correlations[33] were computed between key DCM parameters extracted from the winning model (i.e., the six A matrix connections between the PCC and the other three DMN nodes (mPFC→PCC, PCC→mPFC, LPL→PCC, PCC→LPL, RPL→PCC, PCC→RPL) and the three continuous subclinical symptom scores. Spearman's ρ was selected due to its robustness to outliers and non-normality.

A total of 21 correlations were computed (seven DCM parameters × three subclinical measures). To control the family-wise error rate, Bonferroni correction[33] was applied, yielding a corrected significance threshold of $\alpha = 0.05/21 \approx 0.0024$. Uncorrected associations at $p < 0.05$ were reported as exploratory findings. Raw (non-centred) symptom scores were used for the correlation analysis.

### RESULTS

#### Parametric Empirical Bayes Analysis

Of the 20 participants with a valid winning-model (Model 11), one participant had incomplete subclinical data and was excluded from all second-level analyses, yielding a final sample of N = 19 (11 male, 8 female) for the PEB and correlation analyses.

#### Full model: Biological sex and subclinical symptom severity

A second-level PEB model was estimated on the A-matrix (intrinsic connectivity) and C-matrix (driving input) parameters of Model 11. Following model reduction, no DCM parameter showed strong evidence (posterior probability, $Pp > 0.95$) for an effect of biological sex, severity of psychotic experiences, or severity of subclinical levels of depression (Supplemental Supplemental Figure ). For severity of subclinical levels of anxiety, several A-matrix connections showed posterior probabilities of approximately 0.50, but the corresponding posterior effect sizes were negligible in magnitude and did not approach conventional thresholds for evidence. Overall, there was no clear evidence that current subclinical symptom severity robustly modulated the effective connectivity parameters of the winning DCM.

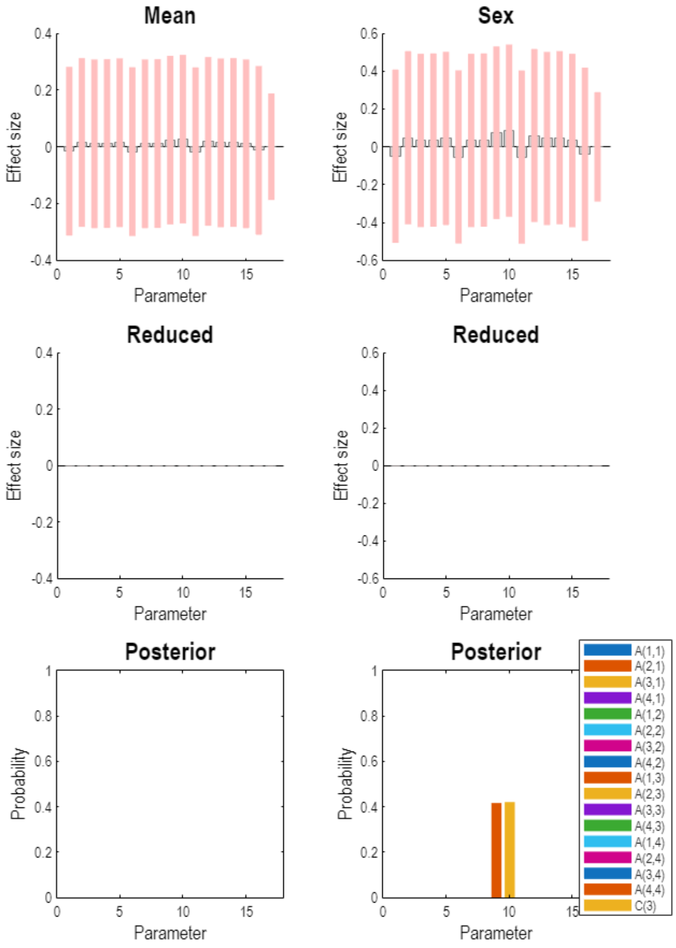


**Supplemental Figure 2. Posterior parameter estimates from the full Parametric Empirical Bayes model (N = 19), shown for the Mean (left column) and Sex (right column) regressors.** Top row: estimated effect size (commonality) for each DCM parameter prior to Bayesian Model Reduction. Middle row: corresponding effect sizes after Bayesian Model Reduction (Reduced); near-zero values across all parameters indicate that the reduction procedure did not retain evidence for these effects. Bottom row: posterior probability of inclusion for each parameter following Bayesian Model Averaging; no parameter exceeded the 0.95 threshold for either regressor.

Post-hoc model: Biological sex only

Given the *a priori* interest in biological sex differences, a post-hoc PEB analysis was conducted. This analysis did not identify evidence for an effect of biological sex on effective connectivity: no parameter reached a liberal posterior probability threshold of 0.49, and BMR did not retain a biological sex-related effect for any connection. This null result is consistent with the absence of a biological sex effect observed in the full PEB model reported above.

Taken together, the PEB and correlation analyses (refer to supplementary materials for correlation analysis results) converge on a consistent pattern: within this proof-of-concept sample, biological sex, and severity of psychotic experiences and subclinical levels of depression and anxiety) were not robustly associated with individual differences in effective connectivity of the winning DCM (Model 11).
